# Endogenous and environmentally modulated leaf shape plasticity in *Boquila trifoliolata*: how good is the evidence for mimicry?

**DOI:** 10.64898/2026.09.03.749126

**Authors:** Fatima Cvrčková, Josef Šonka, Radek Bezvoda, Jana Krtková, Hana Konrádová, Viktor Žárský

## Abstract

Reports that the South American climbing vine *Boquila trifoliolata* modifies its leaf shape to mimic leaves of its host plant have raised considerable attention. Boquila leaf development was proposed to reflect visual inputs perceived in an unknown manner. However, the very existence of Boquila leaf shape mimicry remains inconclusive, since the reported observations are open to alternative explanations. We performed an initial quantitative characterization of leaf shape variability in glasshouse-cultured clonal Boquila plants grown without close host contact, and attempted to reproduce the leaf shape mimicry phenomenon under controlled culture conditions using live olive (*Olea europaea*) plants, artificial (decoy) olive or artificial ivy plants as supports. While we found leaves of variable shape (including those resembling ivy leaves) both in plants grown without host contact and in those climbing on live or decoy hosts, we did not observe any link between leaf shape and host presence or character. Instead, the leaf shape appears to be affected by environmental conditions including the season. Although our study was limited to a single Boquila genotype and an ecologically unrealistic host selection, our observations suggest that the alleged mimicry might originate from unintentional misinterpretation of intrinsic discontinuous leaf shape plasticity in field observations.

## Introduction

*Boquila trifoliolata* (DC.) Decne., the single representative of a monotypic genus of the Lardizabalaceae family (Ranunculales), is a perennial woody twinning wine endemic to evergreen and semi-deciduous temperate forests of Chile and Argentina (Christenhusz, 2012). This plant received little scientific attention outside of systematic botany or ecology literature (see, e.g., Carrasco-Urra and Gianoli, 2009) until a 2014 report of Boquila leaves mimicking the shape of its hosts’ leaves in the wild (Gianoli and Carrasco-Urra, 2014). This observation, reminiscent of previously reported epiphyte-host cryptic leaf shape mimicry in Australian and New Zealand mistletoes (Barlow and Wiens, 1977; Pannell, 2014), made Boquila an instant plant celebrity (e.g., Akpan, 2014; Fiorello et al., 2020) despite the absence of a generally accepted causal hypothesis regarding the underlying ecological, developmental or signalling processes.

Proposed explanations of the Boquila mimicry phenomenon, which can be understood as environmentally induced leaf shape plasticity (Pannell, 2014), include possible response to volatiles from the host, horizontal gene transfer, or response to signals from shared microbiome (Gianoli and Carrasco-Urra, 2014; Gianoli, 2017; Gianoli et al., 2021). However, it has been also proposed that Boquila might be able to sense and perceive the shape of the host leaves by means of a vision-like mechanism, employing hypothetical ocelli – plant-specific structures capable of directional light sensing (Baluška and Mancuso, 2016; Mancuso and Baluška, 2017; Yamashita and Baluška, 2022). This hypothesis inspired an experimental study examining the effects of an artificial host on Boquila leaf shape that revealed statistically significant differences between leaves grown close to an artificial plant compared to those on distant parts of the vine, which was interpreted as consistent with a vision-like perception of host leaf shape (White and Yamashita, 2022).

Although Boquila leaf mimicry is widely accepted as a scientifically established fact (e.g. Segundo-Ortin and Calvo, 2021; Gorelick, 2024), the reported primary evidence for this phenomenon is mostly qualitative and based on limited data from field observations. Published image data consist of photos of a total of 19 individual Boquila – host pairs, with the degree of host – vine leaf shape similarity somewhat varying (Gianoli and Carrasco-Urra, 2014; Gianoli, 2017; Gianoli et al., 2021). The only experimental study of Boquila mimicry so far (White and Yamashita, 2022) was initiated as a citizen scientist project where control and host-exposed (mimic) leaves were collected from branches of different ages that probably experienced substantially different light conditions (see Yamashita de Olivieira, 2024). Its conclusions therefore should be interpreted cautiously.

In this report, we are establishing a methodological framework for investigating the Boquila leaf shape plasticity and its presumed relationship to host leaf shape. This approach, which is readily adaptable for field studies, is then applied to characterize endogenous leaf shape plasticity in glasshouse-grown Boquila plants. Lastly, we are attempting to reproduce the leaf shape mimicry phenomenon under controlled conditions, including quantitative evaluation of host-Boquila leaf shape similarity. Our observations indicate the presence of intrinsic discontinuous Boquila leaf shape variability that might have been unintentionally misinterpreted as mimicry in field observations.

## Results

### Boquila leaf shape is intrinsically variable

All our experiments employed clonal Boquila plants derived from an individual growing in the glasshouse of the Teplice botany garden since 2012. For gaining an initial insight into the leaf shape variability, we harvested all healthy leaves from ten terminal shoots of solitary Boquila clones growing in our experimental glasshouse in Prague (for an example shoot see Figure 1A). Leaf collection took place between September 2024 and March 2025, from shoots that grew in late spring, summer, fall and winter. In addition, we collected both terminal shoots and loose randomly selected leaves from older stems of the parental plant in Teplice, which was climbing a man-made support without direct contact with heterospecific hosts (Figure 1B), in January 2026. In this case, we kept separate leaf pools from the top, middle and bottom thirds of the canopy height in order to detect possible differences between leaves grown at different light levels (and possibly also times, though we had no possibility to estimate the age of the sampled branches). Undamaged leaves from the Teplice sample have been also used for a detailed analysis of the endogenous Boquila leaf shape variability to be reported elsewhere (Neustupa et al., *in preparation*).

**Figure 1.**
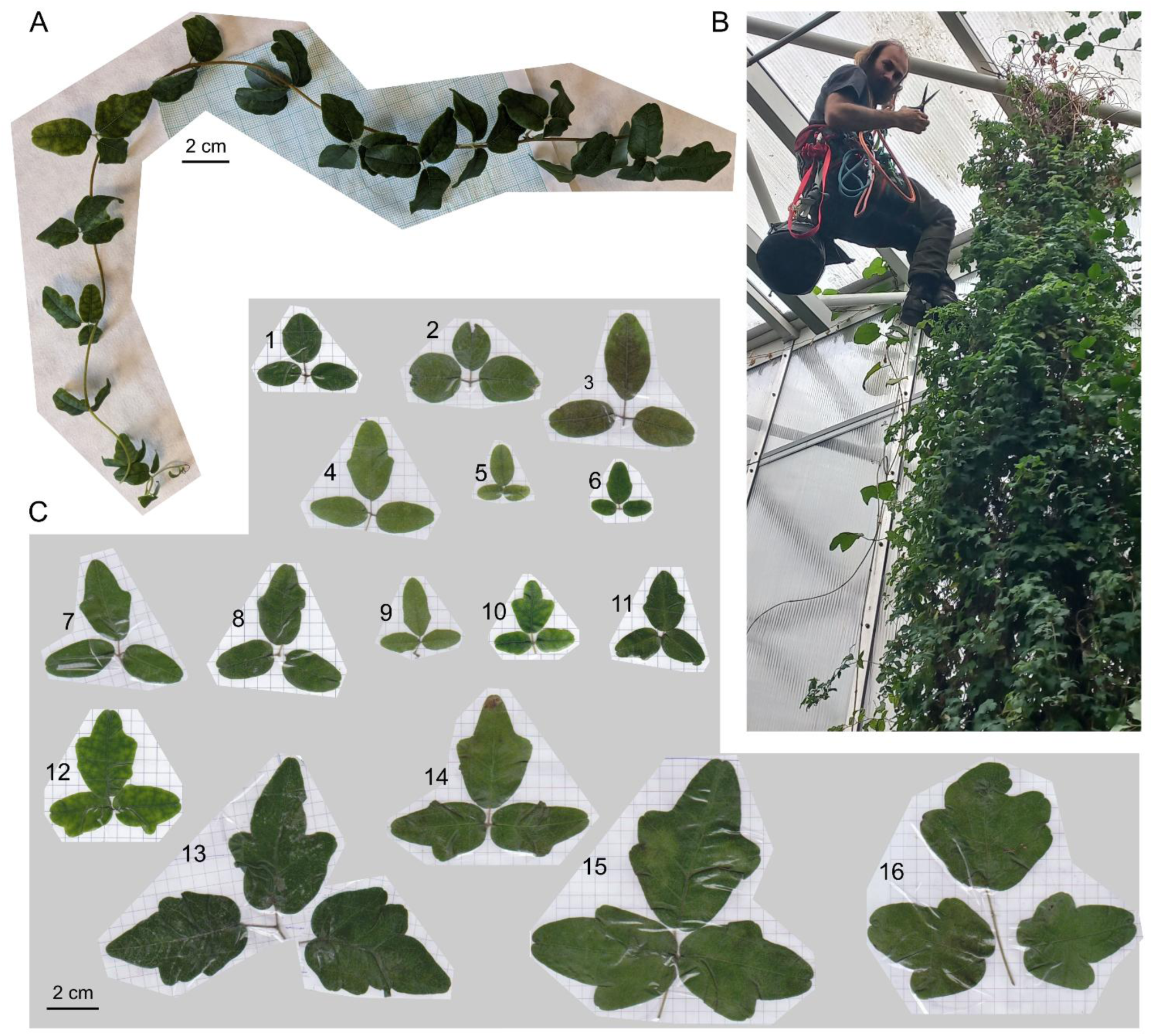
Examples of Boquila leaf shape variability. (A) A shoot harvested from a solitary plant in the Prague glasshouse. (B) Co-author Josef Šonka collecting samples from top of the canopy in the Teplice glasshouse. (C) Examples of leaf shapes from plants grown without direct contact with a plant host. For leaf identification in deposited images and measurement data tables see Supplementary Table S1.

Among the collected leaves, we noticed a substantial variability of both size and shape (Figure 1C; see also Supplementary Table S1). Overall shape of individual leaflets ranged from nearly isodiametric (e.g. leaves 1, 2 and 16 in Figure 1C) to very elongated (leaves 4, 7 and 9). Leaflets also exhibited varying numbers of lobes and indentations, from zero (entire margins, e.g. leaves 1, 2, 3, 5) to six (bottom leaflets of leaf 13). If at least one of the leaflets was lobed, the lobe number was typically higher in the central leaflet than in the side ones (leaves 4, 6-12, 15), although in some leaves these numbers were identical (leaves 14 and 16). While the central leaf was usually symmetrical with rare exceptions (leaf 15), lateral leaflets sometimes exhibited an asymmetric shape with lobes less prominent or absent on the side adjacent to the central leaflet (leaves 11, 12 and 14).

This leaf shape plasticity, comparable to or exceeding the shape range documented in the leaf mimicry reports ((Gianoli and Carrasco-Urra, 2014; Gianoli, 2017; Gianoli et al., 2021) was obviously due to intrinsic (or abiotic environmental) causes, since we took care to sample branches and leaves at the distance of more than 25 cm from individuals of other plant species (i.e. from shoots growing loosely or twinning either on inanimate supports or on Boquila itself) to exclude possible influences of live plant hosts.

### Defining parameters for quantitative description of Boquila leaf morphospace

For quantitative description and subsequent analysis of the Boquila leaf morphospace, we initially selected a set of 26 quantitative leaf parameters that were amenable to semi-high-throughput measurements (Table 1; see also Materials and Methods). Briefly, each leaf was treated as a unit consisting of three leaflets, each of them characterized by eight parameters: area, perimeter, standard ImageJ shape descriptors (circularity, aspect ratio, roundness and solidity – see Ferreira and Rasband, 2012), rectangularity and form factor (White and Yamashita, 2022). Two additional parameters characterized the leaf as a whole – a “central ratio”, or the central leaflet size expressed as a fraction of the total leaf area, and an “asymmetry factor” capturing the area difference between left and right leaflets normalized to the central leaflet area.

**Table 1:** Quantitative parameters used to characterize Boquila leaf shape.

| **Parameter** | **Unit** | **Definition** | **Core** |
| --- | --- | --- | --- |
| **C_Area** | mm<sup>2</sup> | Central leaflet area | no |
| **C_Perim** | mm | Central leaflet perimeter | no |
| **C_Circ** | dimensionless | Central leaflet circularity<br>($4\pi \cdot \text{area} / \text{perimeter}^2$) | yes |
| **C_AR** | dimensionless | Central leaflet aspect ratio<br>(major/minor axis ratio of bounding ellipse) | yes |
| **C_Round** | dimensionless | Central leaflet roundness<br>($4 \cdot \text{area} / (\pi \cdot \text{major\_axis}^2)$) | no |
| **C_Solidity** | dimensionless | Central leaflet solidity (area/convex hull area ratio) | yes |
| **C_Rectangularity** | dimensionless | Central leaflet rectangularity<br>(bounding rectangle area/leaf area) | yes |
| **C_Form_factor** | dimensionless | Central leaflet form factor<br>(width/height of bounding rectangle) | no |
| **L_Area** | mm<sup>2</sup> | Left leaflet area | no |
| **L_Perim** | mm | Left leaflet perimeter | no |
| **L_Circ** | dimensionless | Left leaflet circularity | yes |
| **L_AR** | dimensionless | Left leaflet aspect ratio | yes |
| **L_Round** | dimensionless | Left leaflet roundness | no |
| **L_Solidity** | dimensionless | Left leaflet solidity | yes |
| **L_Rectangularity** | dimensionless | Left leaflet rectangularity | yes |
| **L_Form_factor** | dimensionless | Left leaflet form factor | no |
| **R_Area** | mm<sup>2</sup> | Right leaflet area | no |
| **R_Perim** | mm | Right leaflet perimeter | no |
| **R_Circ** | dimensionless | Right leaflet circularity | yes |
| **R_AR** | dimensionless | Right leaflet aspect ratio | yes |
| **R_Round** | dimensionless | Right leaflet roundness | no |
| **R_Solidity** | dimensionless | Right leaflet solidity | yes |
| **R_Rectangularity** | dimensionless | Right leaflet rectangularity | yes |
| **R_Form_factor** | dimensionless | Right leaflet form factor | no |
| **Central_Ratio** | dimensionless | $C\_Area / (C\_Area + L\_Area + R\_Area)$ | yes |
| **Asymmetry_Factor** | dimensionless | Absolute value of $(L\_Area - R\_Area) / C\_area$ | yes |

To gain initial insight into the Boquila leaf morphospace, we measured 98 leaves from the solitary clonal plants from the Prague glasshouse (Supplementary Table S2) and performed principal component analysis (PCA) to reduce data dimensionality. The results revealed a continuous data distribution with some outliers, with nearly 98 % of the variability accounted for by the first principal component (PC1), which, in turn, was exclusively dependent on parameters reflecting leaf size (Supplementary Figure S1). This indicated that the differences in area and perimeter of individual leaflets would effectively obscure any variability carried by the remaining parameters. These size-dependent variables thus should be excluded from subsequent analyses.

Furthermore, analysis of the mutual correlation between all parameter pairs on the same dataset (Supplementary Figure S2, Supplementary Table S2) revealed very strong and highly significant correlation (and therefore redundancy) not only between each leaflets’ area and perimeter, but also between the aspect ratio, roundness (both reflecting features of the leaflet bounding ellipse) and form factor (a descriptor of the leaflet bounding rectangle). We thus decided to reduce the set of parameters to avoid redundancy between interdependent parameters, resulting in the definition of 14 core leaf parameters that were used in all subsequent analyses (see Table 1).

We next measured the core parameters of over 250 leaves from the parent (Teplice) plant and examined the distribution and mutual correlation of core parameters for 337 leaves from both Prague and Teplice sets combined. This dataset (Supplementary table S3) will be further referred to as the baseline dataset. All parameters exhibited continuous (in some cases markedly asymmetric) distribution with a single maximum, and a varying degree of mutual correlation, with strongest correlations found between mutually corresponding parameters of the individual leaflets (Supplementary Figure S3, Supplementary Table S3). Principal component analysis of the baseline dataset again indicated a continuous data distribution with some outliers, with approximately 70 % of the variability captured by the first three principal components (Figure 2A). Loading of the first three principal components was distributed between the parameters, with PC1 mainly reflecting aspect ratios of the three leaflets and PC2 reflecting multiple variables with a prominent role of the asymmetry factor (Figure 2B).

**Figure 2.**
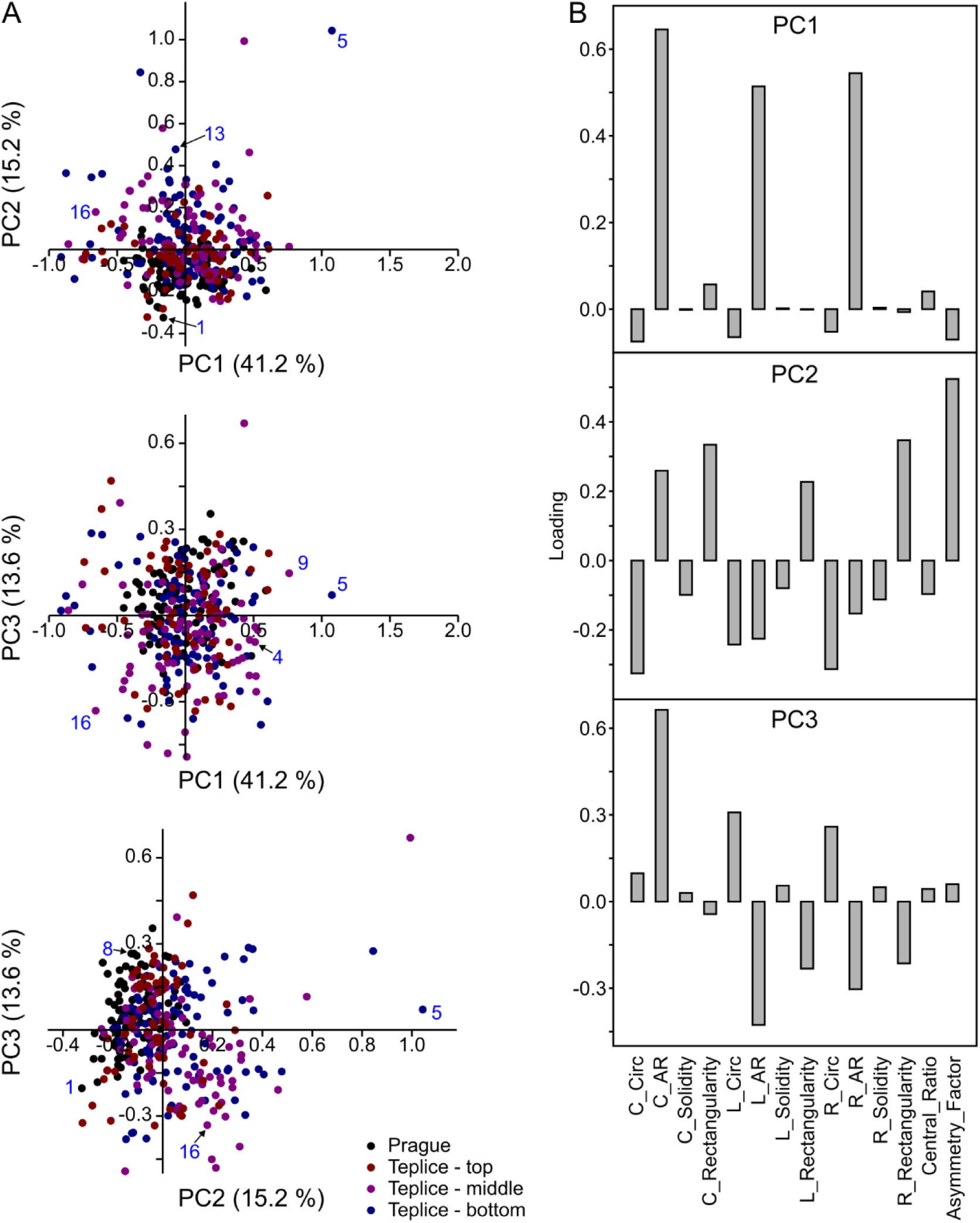
Baseline Boquila leaf shape variability. (A) Scatterplots of PCA results calculated from the core parameters set of 337 leaves from solitary plants from the Prague and Teplice glasshouses (points correspond to individual leaves, color-coded by sample source; selected individual leaves from Fig. 1C are labelled by numbers in blue). Fraction of variance attributable to each PC is shown in parentheses. Only leaves with data for all three leaflets available were considered. (B) Loadings of the first three principal components. For source data, see Supplementary Table S3.

### Boquila can produce at least two distinct leaf categories

While PCA did not reveal any obvious patterns or clusters in the multidimensional baseline data, this method is primarily performing a linear dimensionality reduction without any attempt at cluster detection. We therefore next applied the uniform manifold approximation and projection (UMAP, Healy and McInnes, 2024) algorithm, which performs a non-linear dimensionality reduction combined with clustering based on local distances between data points, on the same baseline data set. We also included into this analysis core parameters measured from a sample of live olive (*Olea europaea* L.), artificial olive and artificial ivy leaves (Figure 3A, Supplementary Table S4) that were to be used as supports in subsequent attempts to induce mimetic leaf response, in order to assess the relationship between baseline Boquila leaf shape variability and host leaf shape.

**Figure 3.**
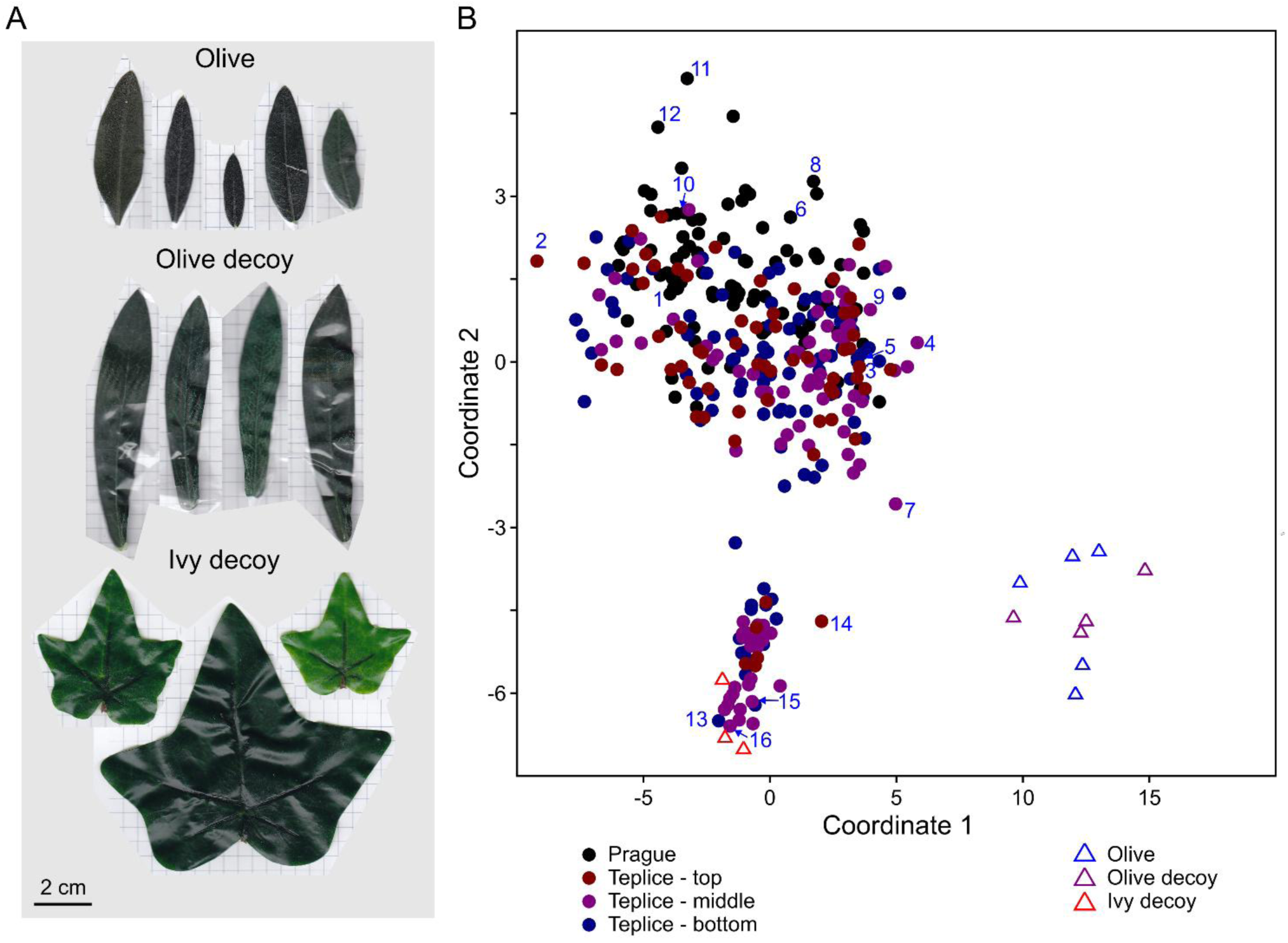
Comparison of baseline Boquila leaf shape variability and host leaf shape distribution. (A) Representative leaves of the live host (olive), host decoy (artificial olive) and arbitrary decoy (artificial ivy). (B) UMAP clustering of 337 leaves from solitary Boquila plants from the Prague and Teplice glasshouses based on core morphometric parameters (the baseline dataset). Points corresponding to individual leaves are color-coded according to the sample source; triangles denote live or decoy host leaves analysed in the same manner. Individual leaves from Fig. 1C are highlighted. For source data, see Supplementary Tables S4 and S5.

Results of this analysis (Figure 3B) indicated the presence of two distinct Boquila leaf populations, both containing a mixture of leaves from the Prague plants and from all three sampled portion of the Teplice canopy. The larger group comprises leaves of relatively simple, less lobed shape (such as leaves 1-12 from Figure 1C), while the other one includes the more prominently lobed ones (leaves 13-16 from Figure 1C) together with the artificial ivy leaves. Notably, olive leaves, both live ones the and quite life-like artificial ones, clustered together but apart from Boquila leaves, confirming that the UMAP algorithm can pick up features corresponding to human-detectable leaf shape similarity.

### An experimental setup to study host effects on leaf shape

To examine the effects of live or artificial hosts on Boquila leaf shape, we set up parallel Boquila cultivations on four different supports – control (C), live host (H), artificial host decoy (HD) and artificial arbitrary decoy (AD) – and collected leaves from shoots grown for several months on these supports. The cultivation took place between August 2024 and May 2026, with three rounds of sampling (in the spring of 2025, in October 2025 and in late May 2026). Because of plant loss in the early spring of 2025 (see below), we replaced about half of the plants in the summer of 2025. We will be referring to the individual sampling rounds as experiment 1 (the early spring 2025 sampling) and experiment 2 (set up in the summer of 2025 with slightly modified growth conditions, see below) with two samplings – in the fall of 2025 and late spring of 2026 (Figure 4A).

**Figure 4.**
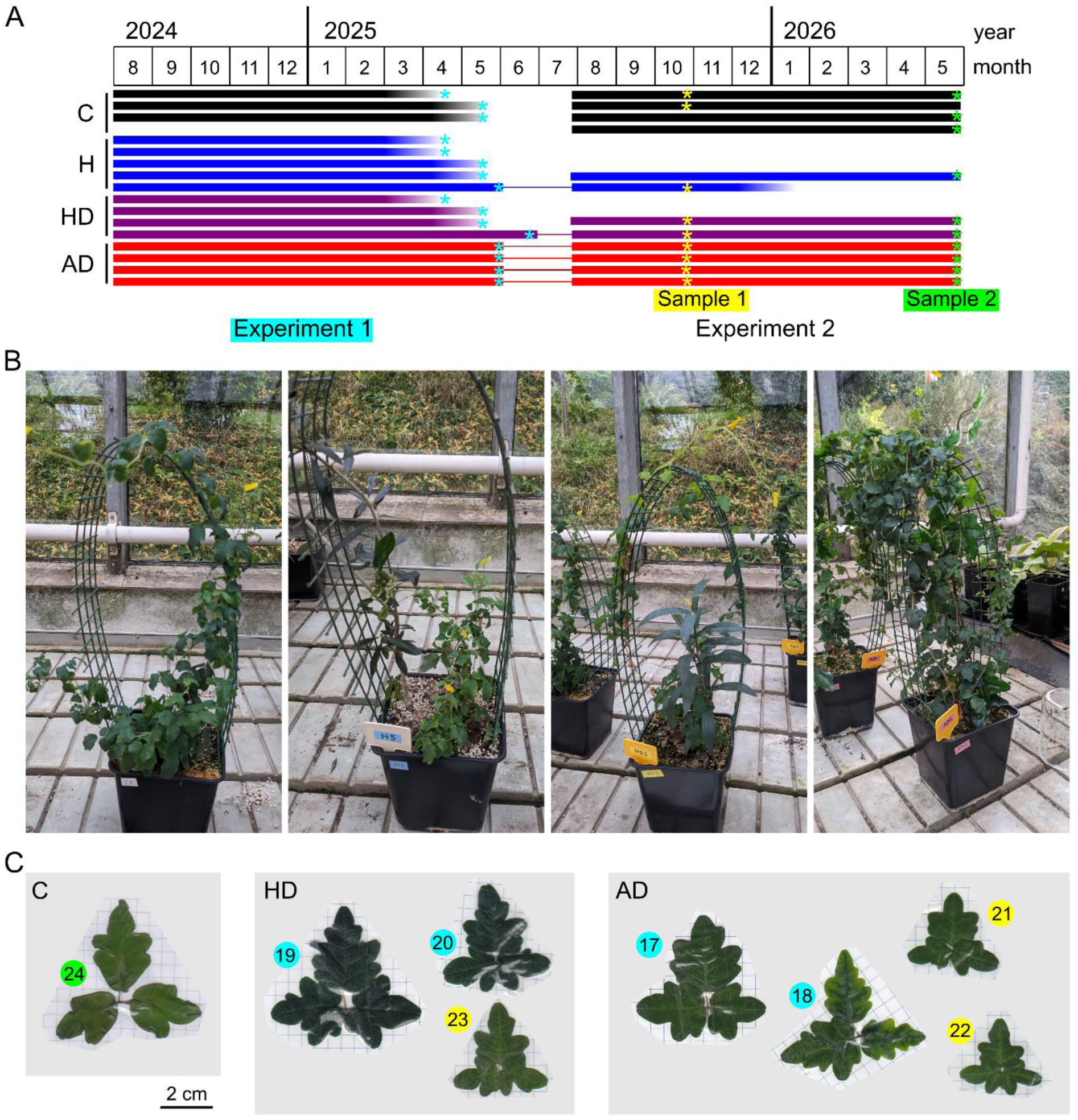
Experimental setup used to study possible host effects on Boquila leaf shape. (A) Schematic of the experiment timeline. Bars correspond to individual plant growth on supports, faded bars denote visibly stressed (chlorotic) or dying plants. Times of sample collection are marked by asterisks, color-coded by experiment and sampling round. Thin lines connecting the two experiments indicate reuse of individual plants. Abbreviations denote the supports: C – control (wire mesh), H – live host (olive), HD – host decoy (artificial olive), AD – arbitrary decoy (artificial ivy). (B) Experimental plants in the glasshouse. (C) Complex-shaped leaves occasionally found in some samplings regardless of the host (hosts denoted by abbreviations, sampling rounds colour-coded as in (A). For leaf identification in deposited images and measurement data tables see Supplementary Table S1.

The culture conditions were as follows. For controls (C), the plants were provided only a wire trellis as a support. For the live host (H) set, the vine was sharing the pot with a young olive tree and a wire trellis. For the decoy host cultures, artificial olive branches (HD) or artificial ivy (AD), made of plastic, wire and textile, were stuck into the substrate in the pot and attached to the trellis (Figure 4B).

In the first experiment, the plants grew vigorously through the fall but slowed down during the winter and most of them started to show signs of stress and damage (chlorosis, leaf death, and in some cases ultimately whole plant death) in the early spring. These plants have thus been sampled prematurely (see Figure 4A). We suspect that the damage may have been triggered by the combination of a short daylength, relatively low humidity and high daytime light intensity (due to the absence of shading by deciduous trees adjacent to the glasshouse), because we noticed that Boquila plants propagated in a shadier part of the glasshouse, as well as the AD plants that were somewhat shaded by the artificial ivy, appeared noticeably healthier. The second experiment has therefore been set up in a part of the glasshouse that receives less direct light in the winter, and all plants but one survived without obvious problems.

During sample collection in some experiments, we noticed the presence of very prominently lobed leaves (Figure 4C). Although it would be tempting to interpret these leaves as “ivy-like”, they occurred on both AD and HD plants, and rarely also on controls, mostly in the fall/early spring samplings (suggesting that they developed from primordia established during the summer or early fall).

### Evidence for host-independent but not host-induced leaf shape variation

Next, we quantified the core morphometric parameters from all harvested leaves (Supplementary Table S5) and subjected them to UMAP analysis in the same manner as the baseline data. In two of the three samplings – Experiment 1 and Experiment 2 sample 1 – we obtained similar discontinuous leaf type distribution as in the previous baseline analysis (Figure 5A, left and centre). Notably, the cluster harboring the artificial ivy leaves in both cases contained leaves from three or all four experimental variants, indicating that increased leaf lobing was not induced by a lobed (artificial) host. In the third sample (Experiment 2 sample 2), which comprised leaves from vigorously growing spring shoots, there were very few “ivy-like” leaves and UMAP did not indicate any obvious clustering, but rather a continuum of leaf shapes with the artificial ivy mapping to one corner, together with more prominently lobed Boquila leaves. Notably, this sample also involved some leaves from shoots that grew prostrate on the glasshouse table or floor rather than twining around any support (“creepers”). These mapped as dispersed among all other control or experimental leaves (Figure 5A, right).

**Figure 5.**
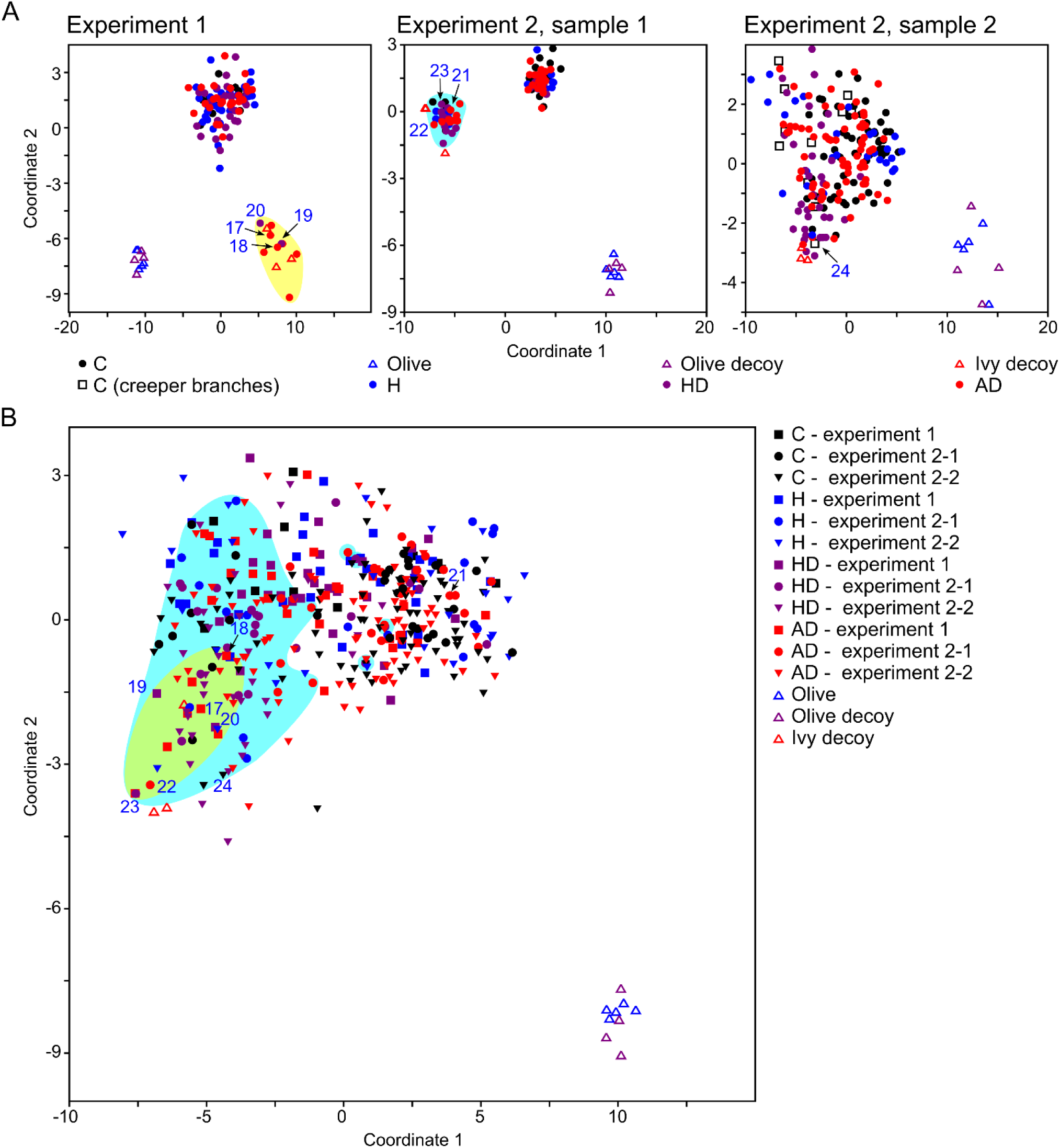
Leaf shape distribution in Boquila grown on various supports. (A) Core parameter-based UMAP clustering performed separately for individual biological replicates and samplings of the host effects experiment, with leaves shown in Fig.4C highlighted. (B) Core parameter-based UMAP clustering of all data from the host effects experiments, pooled together. Yellow and cyan areas mark the leaves that clustered together in experiment 1 and in sample 1 of experiment 2. For source data, see supplementary tables S4 and S5.

A similar pattern of a continuous leaf shape distribution was also observed upon combination of all three samplings together (Figure 5B), or when the data from the host effect experiments were combined with the baseline data (Supplementary Figure 4A). Notably, in the latter case, the previously documented two baseline leaf categories (compare Figure 3B) were no longer apparent, suggesting that the addition of large number of leaves from spring-grown branches (Experiment 2 sample 2) reconfigured the UMAP – generated morphospace map, resulting in disappearance of the two clusters. Consistent with this interpretation, UMAP on a combination of baseline data and all controls from all experiments also produced a continuous pattern rather than a discontinuous one (Supplementary Figure 4B).

In summary, our observations suggest that at least at some growth conditions Boquila can produce very diverse leaf shapes regardless of the presence or properties of its host, although the extent and character of leaf plasticity may be affected by other factors such as the season.

## Discussion

The reports of cryptic leaf shape mimicry in *B. trifoliolata* gained considerable and lasting attention not only inside but also outside the fields of plant sciences and ecology (see Introduction; also, e.g., Jones, 2022; Schlanger, 2024; Bourdev, 2025; García, 2025). Boquila mimicry has even been used as an argument in favour of “intelligent design” (ÓLeary, 2024). Yet, at the same time, the scientific community remains somewhat uncomfortable with this phenomenon, and possibly with the idea of leaf mimicry in general. For example, a short review on plant mimicry written at the height of Boquilla’s popularity (Pannell and Farmer, 2016), co-authored by one of the participants in the vigorous discussion that followed the first Boquila mimicry report (Pannell, 2014), does not mention Boquila at all. Even the similarity of leaf shape between Australian mistletoes and their hosts, long considered a prototypic case of cryptic leaf mimicry, has been recently explained as a result of evolutionary convergence driven by similar ecological selection pressure (Midgley, 2026).

One of the reasons for the plant biologists’ discomfort with Boquila mimicry may be the lack of a plausible hypothesis explaining the reported observations (see Introduction). Unlike the mistletoes, whose leaf shape is a species-specific feature established in the course of evolution (Barlow and Wiens, 1977), the similarity between Boquila and host leaves appears to result from developmental leaf shape plasticity, with leaves of the same branch reported to attain different shapes depending on the closest host plant (Gianoli and Carrasco-Urra, 2014). Another cause of concern, however, may be the character of published evidence for the phenomenon, which rests solely on observations from field studies that may be intrinsically prone to unintentional observer bias (compare Chadwick et al., 2024). Narrative descriptions of host – Boquila similarity are accompanied by illustrative photos (Gianoli and Carrasco-Urra, 2014; Gianoli, 2017; Gianoli et al., 2021), but no systematic sampling has been reported and almost no quantitative data are provided. There is only a summary of measurements documenting statistically significant correlation between Boquila and host leaf size, petiole length, angle and colour, i.e. parameters not reflecting leaf shape, together with evidence of differences between prostrate (creeper) and climber shoots in the same parameters (Gianoli and Carrasco-Urra, 2014). Thus, the evidence for leaf shape similarity between the vine and its host is mainly qualitative and documented by limited amount of data, in contrast to well characterized similarity of Boquila and host epi-and endophytic microbiomes (Gianoli et al., 2021).

The only relevant report employing quantitative leaf shape evaluation until now is the experimental study of Boquila development on an artificial host (White and Yamashita, 2022). However, this paper, unfortunately, suffers from some methodological drawbacks that make the reported observations open to alternative explanations. Namely, the “treated” (artificial host-exposed) plants and the controls were separated by an opaque shelf that may have significantly affected the light conditions, and thus the comparison between “mimic” and “non-mimic” leaves could have at the same time been between light and shade grown ones. Moreover, the authors obviously treat individual leaflets as “leaves”, which could have introduced some confusion (the “single lobe mimic leaf” of White and Yamashita’s Figure 3 is very obviously an asymmetrical side leaflet). Nevertheless, we credit this report for inspiring the design of our present study, where we attempted to reproduce its results in a better controlled manner and, at the same time, gain an insight into the background, or endogenous, variability of Boquila leaf shape.

While we did not observe any evidence of Boquila mimicking the shape of host leaves (be the host a live plant or an artificial one), we did document endogenous leaf shape plasticity that in some situations resulted in the presence of two distinct leaf morphotypes, including one remarkably reminiscent of the (artificial) ivy that was used as the host. However, the “ivy-like” leaves were produced also by plants that had no contact with the “model”, and therefore they cannot be considered as mimic. We rather suspect that these leaves are the outcome of a natural seasonal leaf shape variation, since they were found nearly exclusively on branches grown in the summer, fall or early spring, but nearly absent on late spring-grown shoots. This is in good agreement with the observation that spring-grown Boquila leaves appear to be narrower and less lobed than summer and winter-grown ones (White and Yamashita, 2022). A recent phylogenetic and modelling study (Malone et al., 2026) indicated that simplification of leaf shape can occur relatively easily by subtle alterations in multiple parameters, resulting in a developmental bias towards simple leaf shape. We can speculate that spring-grown leaves, which develop from primordia initiated in winter under the glasshouse conditions, may be especially prone to this developmental bias.

We need to stress that our present observations, while providing no evidence for leaf mimicry, should not be interpreted as evidence of its absence. Our choice of olive as the host was motivated by the access to live plants with approximately Boquila-like leaf size and commercially available artificial copies. However, the mimetic response (or other host-induced leaf shape alteration) may depend on signals received from the native hosts that have co-evolved with Boquila over a long time. Indeed, there are documented examples of volatiles produced by healthy plants modulating biotic stress responses or even growth of heterospecific or conspecific neighbours (Himanen et al., 2010; Kheam et al., 2024; Åbonde et al., 2026). In the case of Boquila, species-specific host-derived volatiles might induce development of distinct leaf morphotypes.

Moreover, the ability to perform the mimetic response may be genotype-dependent. IOur experiments were conducted on the clonal progeny of a single plant, and in all likelihood, our plants were of a different genetic constitution than those observed by E. Gianoli and co-workers. Indeed, detailed examination of photos in their reports (Gianoli and Carrasco-Urra, 2014; Gianoli, 2017; Gianoli et al., 2021) reveals leaf forms we never observed (especially serrated leaves) but no highly lobed leaves such as those in our Figure 4C. In addition, our source individual has been growing for at least 12 years under stable glasshouse conditions, and it is conceivable that some epigenetic changes affecting its developmental plasticity took place during that time.

In summary, although we failed to reproduce Boquila leaf shape mimicry under controlled experimental conditions, we consider the possibility of this phenomenon existing in the wild open. The simple and accessible methodology for quantitative leaf shape description provided in the present report is based on the use of free software, and, in principle, adoptable to field studies. We encourage colleagues with access to native Boquila habitats to apply our or similar methods in a future rigorous quantitative study of the leaves of Boquila plants growing in their native environment on their native hosts, which would conclusively settle the question of the existence of leaf mimicry in this species.

## Materials and Methods

### Plants

*Boquila trifoliolata* (DC.) Decne. has been clonally propagated from cuttings collected from an individual (growing in the Valdivian Forest section of the Chile flora collection maintained in the Teplice Botany Garden (IPEN Code TEBLI, https://www.botanickateplice.cz/) under the Florius catalogue (http://florius.cz/) code ACCID 2012.11312. Under our glasshouse conditions (see below), the plants grew vigorously during the spring and fall but exhibited frequent wilting, blackening and die-back of young shoots in winter (possibly due to the activity of undiagnosed fungal pathogens) and during hot, dry periods in the summer, especially when temperatures exceeded 30 °C. However, the plants typically recovered well due to activation of axillary buds on healthy mature shoots when conditions became more permissive. In general, the plants were producing two distinct types of shoots – “climbers” right-hand twinning around available supports and prostrate “creepers” (runners, stolons) growing horizontally. Individual shoots were to some extent able to alternate between these modes (Supplementary Figure S5; compare Gianoli and Carrasco-Urra, 2014). The creepers were spontaneously rooting at damp places and producing new climbers from axillary buds. Tips of creepers also sometimes changed their mode of growth to right-hand twinning and became climbers. Tips of climbers that could not find a suitable support usually dried off, followed by production of new shoots from axial buds. Together, this resulted in the plant being able to spread in space quite rapidly and efficiently (compare Figure 1B).

Olive (*Olea europaea* L.) cv. Picual plants have been derived from cuttings produced from long-term *in vitro* cultures (Rejšková et al., 2007). For *ex vitro* acclimatiation, 10 weeks old well-lignified and rooted apical microshoots were carefully harvested from the culture vessels. The remaining medium was gently washed from the root systems with lukewarm distilled water to minimize the risk of subsequent microbial decay. The plantlets were then transplanted into plastic pots containing a wet sterilized substrate mixture of peat and perlite (1:1, v/v). To prevent rapid desiccation, the transplanted plantlets were initially maintained under high relative humidity using transparent covers in a controlled growth chamber. Over a period of 5 weeks, the relative humidity was progressively decreased to ambient levels (approx. 60%). The growth chamber was maintained at a temperature of 24 ± 2 °C under a 16-hour photoperiod with a light intensity of 200 µmol m⁻² s⁻¹ provided by cool-white fluorescent lamps. Following this hardening phase, the acclimated olive plants were transferred to standard greenhouse conditions (see below) for further vegetative development.

Olive plants were subsequently propagated vegetatively as follows. 20-25 cm long cuttings from mature shoots were treated by application of the AS-1 rooting stimulant containing NAA and vitamin B3 in a talcum powder base (AgroBio Opava, CZ) and rooted in a mix of equal amounts of pit sand, perlite and Hawita tray substrate (70 % white peat, 30 % black peat, PG mix starter fertilizer 0.5 kg/m^3^, increased iron contents, 0-5 mm granularity; Sinco, CZ). Cuttings were maintained under 60 - 90 % relative humidity, temperature range 15-25 °C, at a semi-shaded glasshouse shelf. Over 80 % of cuttings established roots within one to two months. Plants were then transferred to a 2:1 mix of the Hawita Basi substrate (white peat, clay 40 kg/m^3^, PG mix 1 kg/m^3^, 10-25 mm granularity; Sinco, CZ) and pit sand with addition of the Osmocote 8-9M fertilizer at 2.5 g/l for further growth and maintenance under natural photoperiod. Temperature range was mostly as above, with occasional winter minima of 10 °C and summer maxima about 35 °C.

### Boquila culture conditions and propagation

Experimental plants were grown in the Charles University, Faculty of Science, Section of Biology experimental glasshouse in 15 x 15 x 15 cm pots under natural daylight on half-shaded shelves, with temperature maintained by ventilation and heating at 15 – 20 °C for vegetative propagation and at 15 – 25 °C for subsequent cultivation and host effect experiments, at relative humidity of 60 – 80 % unless stated otherwise.

For vegetative propagation, mature creeper shoots and small pots (10 x 10 x 10 cm) were typically used. Two propagation protocols were alternatively employed. In the first approach, two-node shoot cuttings treated with the AS-1 rooting stimulant were planted into pots containing a 1:2 mix of Hawita tray substrate and perlite and maintained either under the above-described conditions, or at increased temperature (25 °C) or increased relative humidity (80 %). Even under conditions giving the best rooting efficiency (standard temperature, 80 % humidity) only about 15 % of cuttings rooted successfully. The second approach involved layering of shoots that were attached to the same substrate as used for cuttings by a bent wire at every other node (Supplementary Figure S6). The substrate was maintained constantly moist (in some cases, sphagnum moss was employed to stabilize humidity, but this did not have a marked effect on rooting efficiency). Rooting typically took place in over 90 % of substrate-attached nodes within one month, at which point also new shoots started growing from axillar buds. Subsequently, individual nodes were separated and plantlets transferred to standard culture pots containing a 3:1:1 mix of Hawita tray substrate, perlite and pit sand. Layering was similarly efficient also when using mature climber shoots, which, however, took longer to root.

For investigating possible host effects, individual rooted Boquila plantlets with approximately 10 cm shoots were transferred to standard culture pots with a 3:1:1 mix of Hawita tray substrate, perlite and pit sand, either alone or together with a young olive tree approximately 40 cm tall, and the pots were equipped with a plastic-covered wire trellis. For the decoy treatments, commercially sourced artificial olive or ivy branches, made of plastic, wire and textile (modified by removal of fruit in case of olives), were inserted into the substrate and affixed to the trellis (see Figure 4B). Plants were kept in the glasshouse as described above for the duration of the experiment; due to weather changes, the relative humidity during the experiment varied between 55 – 80 %.

### Leaf sampling and shape quantification

For initial characterization of the Boquila leaf morphospace, whole terminal shoots and loose leaves were collected as described in Results. For the host effect experiments, initial two sampling rounds (Experiment 1 and Experiment 2 sample 1) were performed in a manner minimizing deleterious effects due to defoliation, i.e. randomly selected leaves were collected, attempting to take every 4^th^ to 5^th^ leaf on a selected shoot, as long as the shoot remained adjacent to the studied host, as far as possible. At the same time, conscious effort was taken to represent all leaf shapes encountered on the plant, which might have led to some overrepresentation of extreme shapes. In the third sampling round (Experiment 2 sample 2), whole terminal branches were collected, or, if the plant twinning did not allow this, all leaves from contiguous shoot segments were harvested. Collected leaves were attached to graph paper using a transparent tape, labelled, stored overnight or for a few days under weight to keep them flat, and scanned on an office scanner. Leaves harvested from contiguous shoot segments have been labelled by numbers or lettering in an ascending order starting from the shoot tip (e.g., 2B11 stands for shoot 2B leaf 11, or Bottom3b for the second leaf of shoot 3 from the bottom of canopy); these labels are used in the supplementary tables. The scanned images have been deposited in the BioImage Archive (https://www.ebi.ac.uk/bioimage-archive/) under accession number S-BIAD4056.

Leaf shape parameters were determined using Fiji (Schindelin et al., 2012) version 1.54p as follows. First, upon opening of the first file of a batch, a global scale has been set based on the size of the graph paper grid. For collecting the full set of morphometric parameters, measurements were set to determine area, perimeter, shape descriptors, and bounding rectangle parameters; for collection of core parameters, the perimeter could be omitted. Next, rectangles containing individual leaves (or leaf groups, if in the same orientation) were selected, copied and new images generated from the Fiji internal clipboard. These individual leaf images were converted to binary, obvious optical artifacts were corrected and individual leaflets separated using the brush tool, and images were rotated to position the central leaflet perpendicularly. Central leaflets were identified using the wand tool and measured using the above settings, with results collected into an Excel spreadsheet. The images were rotated again to position the side leaflets vertically and side leaflets were measured in the same manner (in some cases this had to be done separately for the left and right leaflet). A spreadsheet with individual leaves in rows and parameters in columns was assembled and the derived parameters of rectangularity, form factor (if measured), central ratio and asymmetry factor were calculated as defined in Table 1.

## Data analyses

Distribution of individual parameters, their mutual correlations and correlation strengths were determined, and pairwise correlation plots generated, as described previously (Bezvoda et al., 2025).

PCA and UMAP analyses were performed and their results visualised using the PAST software (Hammer et al., 2001) version 5.3, with all variables treated as ordinal. For PCA, the correlation matrix method was used, for UMAP, the Euclidean distance method. Since UMAP cannot handle missing data, any missing values (such as lost or visibly damaged leaflets) were imputed according to the following algorithm: (1) If there is only one leaflet, triplicate it (used especially for the simple-leaved hosts). (2) If a side leaflet is missing, mirror the remaining one. (3) If a central leaflet is missing, mirror the left leaflet.

## Data availability

All leaf images evaluated in this study have been deposited at the BioImage Archive with accession S-BIAD4056, https://doi.org/10.6019/S-BIAD4056. Raw and processed quantitative data used in analyses presented in this study are provided in Supplementary Tables of this manuscript.

## Author contributions

FC – Conceptualization, Formal analysis, Investigation, Writing – original draft preparation, Visualisation; JŠ – Investigation, Writing – review & editing; RB – Formal analysis, Investigation, Data curation, Writing – review & editing; JK – Investigation; HK – Investigation, Writing – review & editing; VŽ – Conceptualization, Writing – review & editing.

## Acknowledgements

We thank Jan Ptáček from the Teplice Botany Garden (IPEN Code TEBLI) for sharing vegetatively propagated *B. trifoliolata* plants and facilitating leaf sample collection. This work has been supported by the V. Kann Rasmussen Foundation grant “Perception and developmental behavior in Boquila: the world according to a mimic vine” to FC.

## Supplementary Materials

Additional File 1 (*.pdf) containing Figures S1 to S6.

Additional File 2 (*xlsx) containing Tables S1 to S5.

