## Additional file 1 for "Endogenous and environmentally modulated leaf shape plasticity in *Boquila trifoliolata*: how good is the evidence for mimicry?"

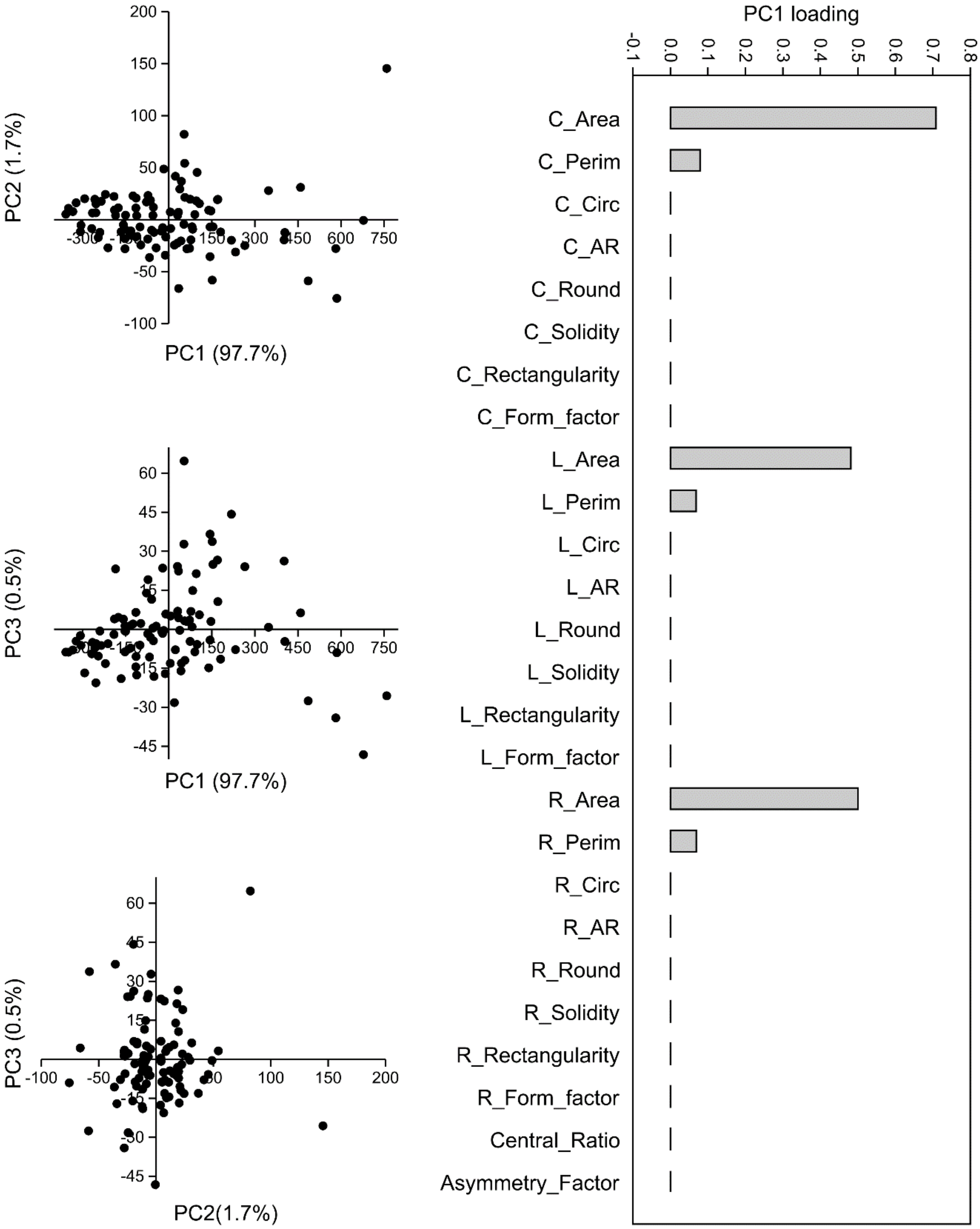


**Figure S1.** Left: Scatterplot of PCA results reflecting the leaf size and shape variability, calculated from all measured morphometric parameters of 98 leaves from solitary plants from the Prague glasshouse (points correspond to individual leaves). Fraction of variance attributable to each PC is shown in parentheses. Right: Loadings of the first principal component. For source data see Supplementary Table S2.

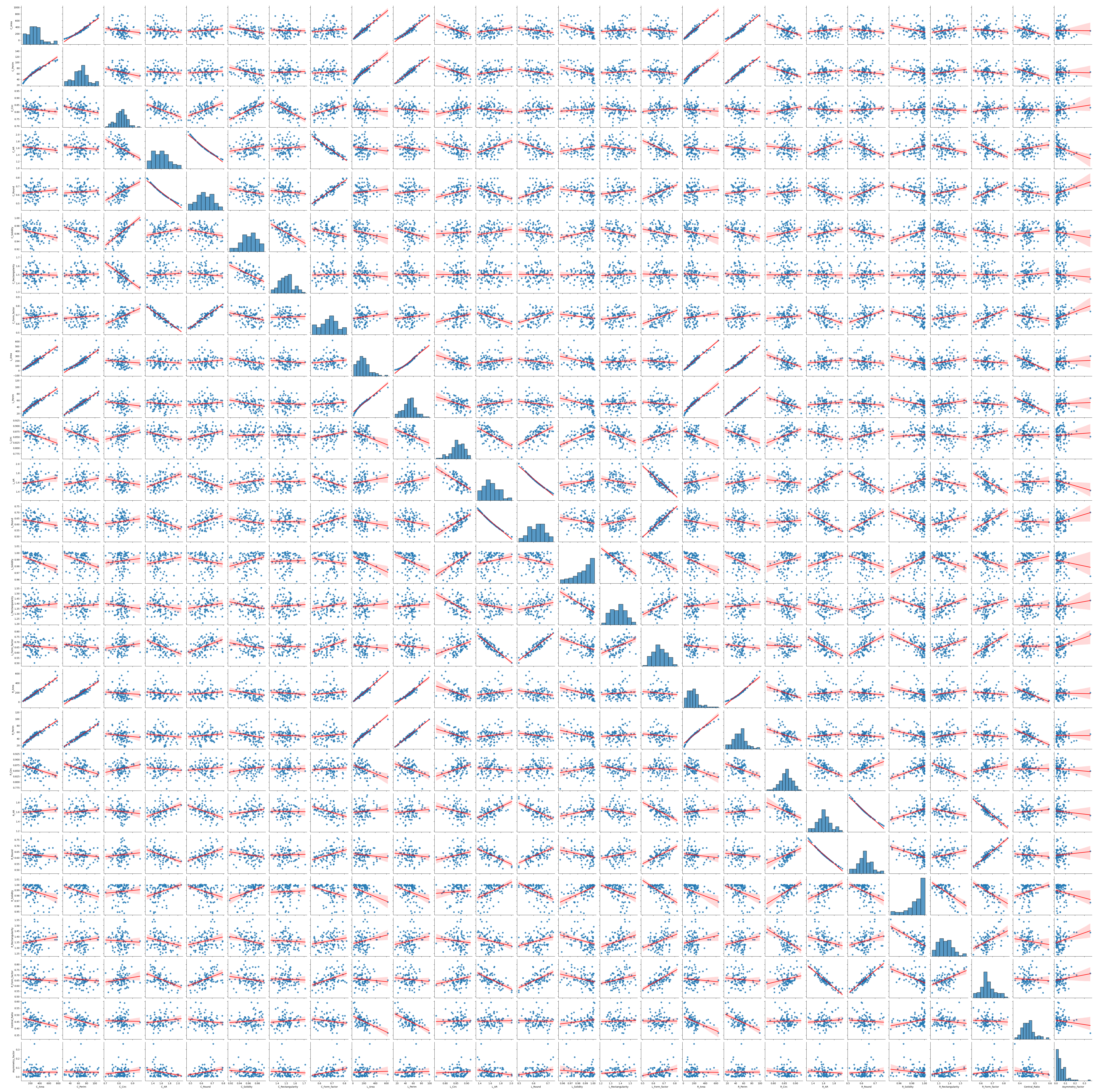


**Figure S2.** Distribution and mutual correlation of all measured morphometric parameters of the leaf set from Figure S1. Distribution of values for each parameter is shown at the diagonal of the matrix, with leaf counts on the Y axis. Source data, Spearman's correlation coefficients (Rho), P-values and coefficient of determination (R2) values for all pairs are provided in Supplementary Table S2.

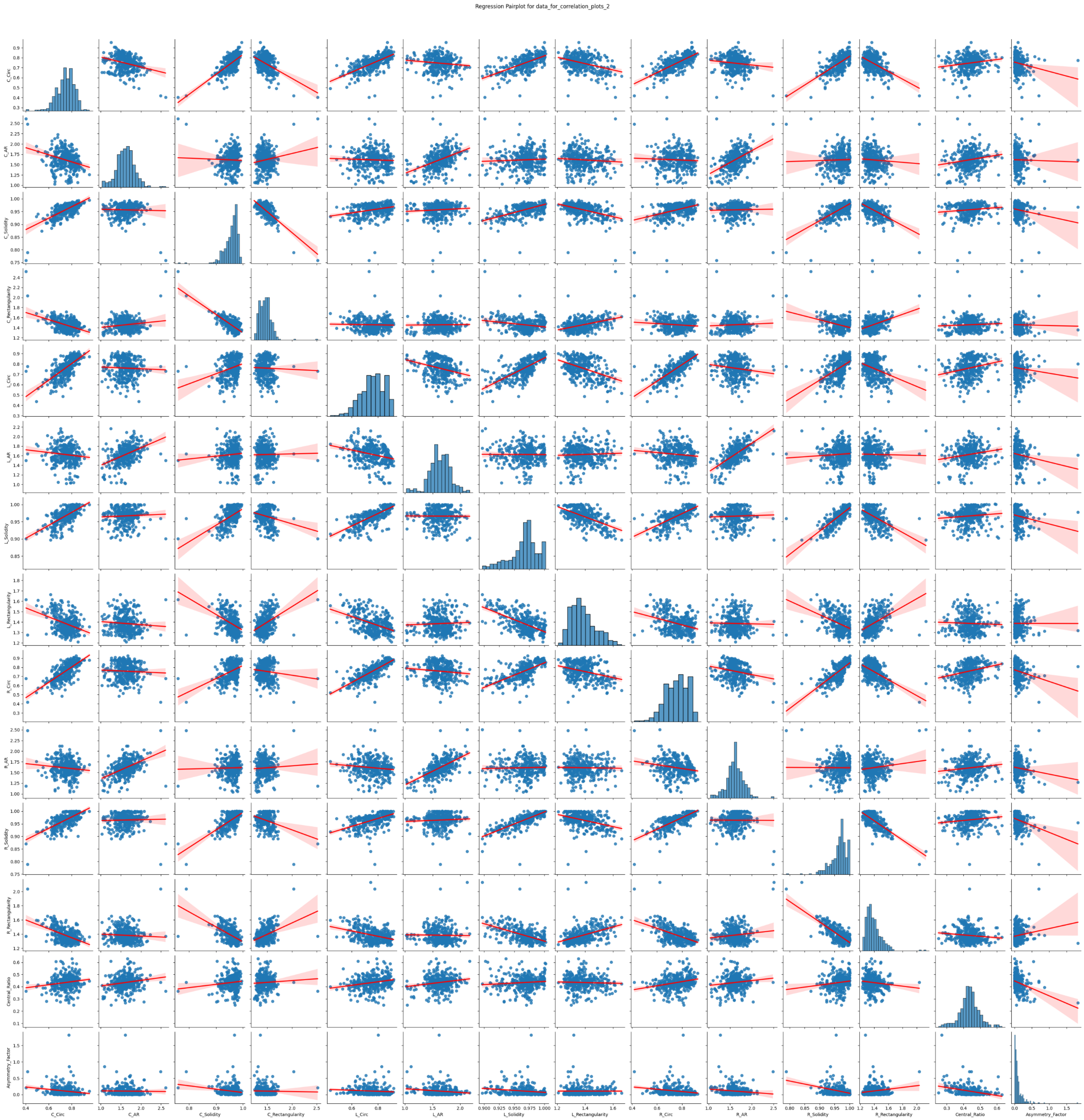


**Figure S3.** Distribution and mutual correlation of core (i.e. mutually independent and size-independent) quantitative leaf shape parameters of 337 leaves from solitary plants from the Prague and Teplice glasshouses, with regression lines and confidence intervals indicated. Distribution of values (on an arbitrary scale) is shown for each parameter at the diagonal of the matrix, with leaf counts on the y axis. For source data, Spearman's correlation coefficients, P values and coefficient of determination ($R^2$) values see Supplementary Table S3.

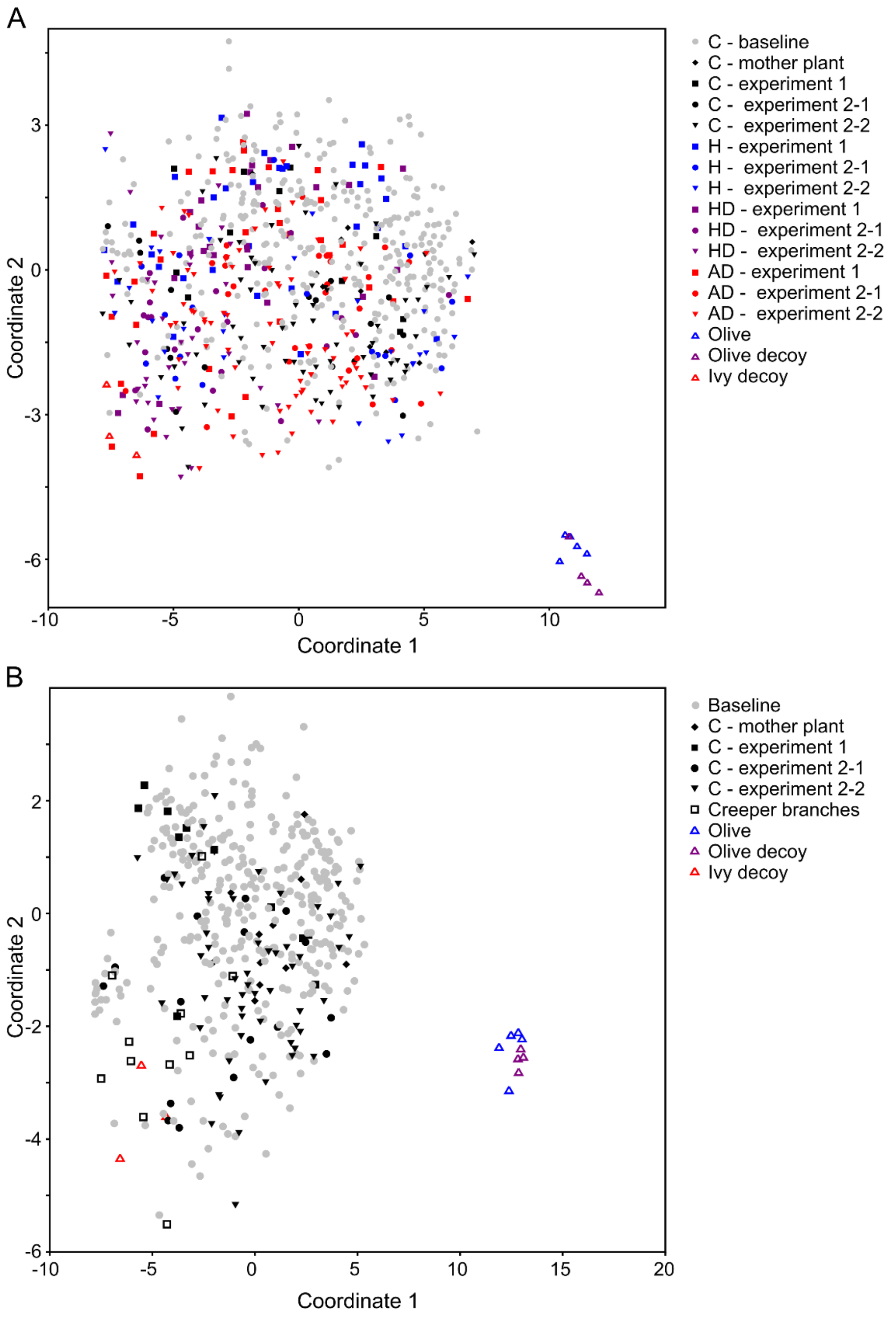


**Figure S4.** (A) Core parameter-based UMAP clustering of all data from the host effects experiments, combined with baseline shape distribution data from both Prague and Teplice glasshouses. (B) Core parameter-based UMAP clustering of control plant leaf data, combining baseline shape distribution data from both Prague and Teplice glasshouses and control leaf data from the host effect experiments. For source data, see supplementary tables S3, S4 and S5.

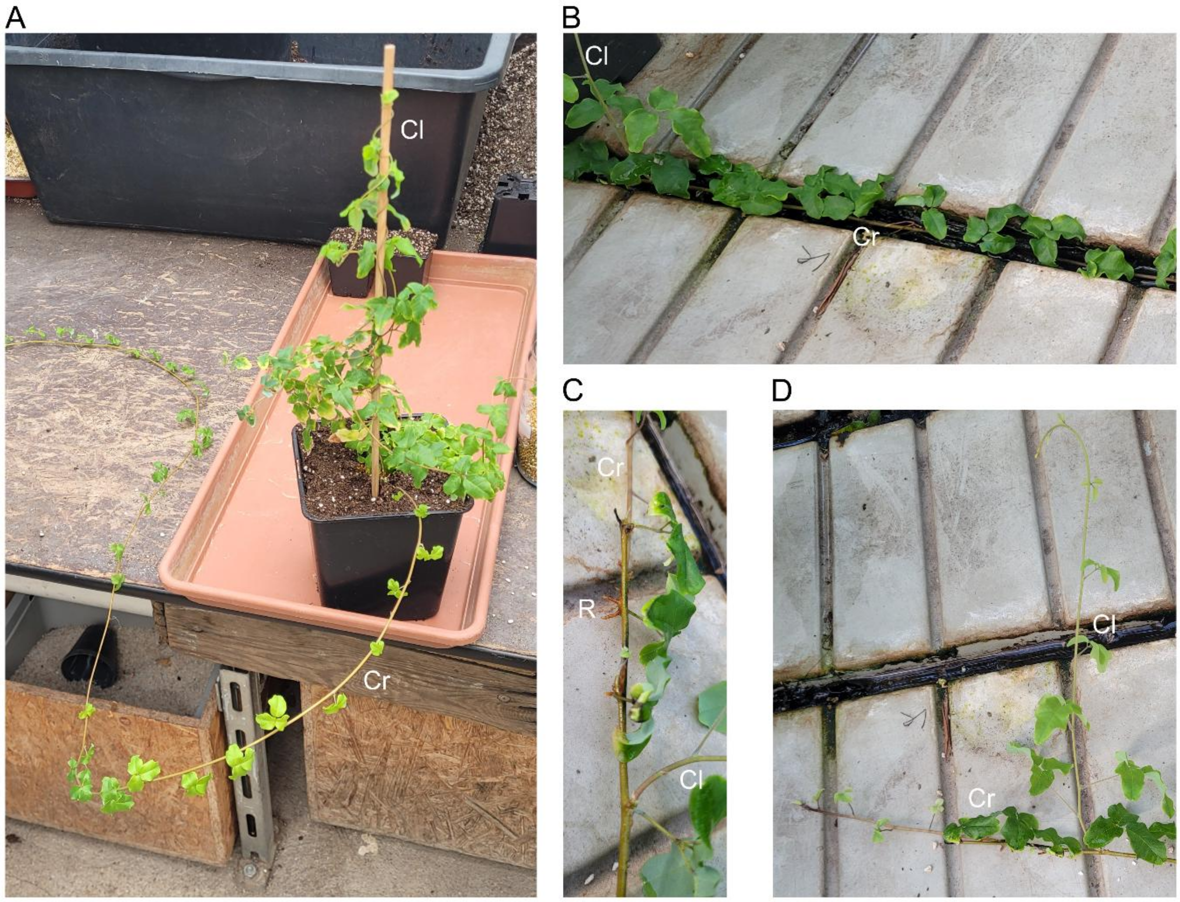


**Figure S5.** Examples of Boquila shoot morphology. (A) The two typical growth modes. (B) A creeper shoot growing along the glasshouse table. (C) Spontaneous rooting of a creeper shoot. (D) A climber shoot growing out of am axillary bud of a creeper. Cr – creeper, Cl – climber, R – adventitious roots.

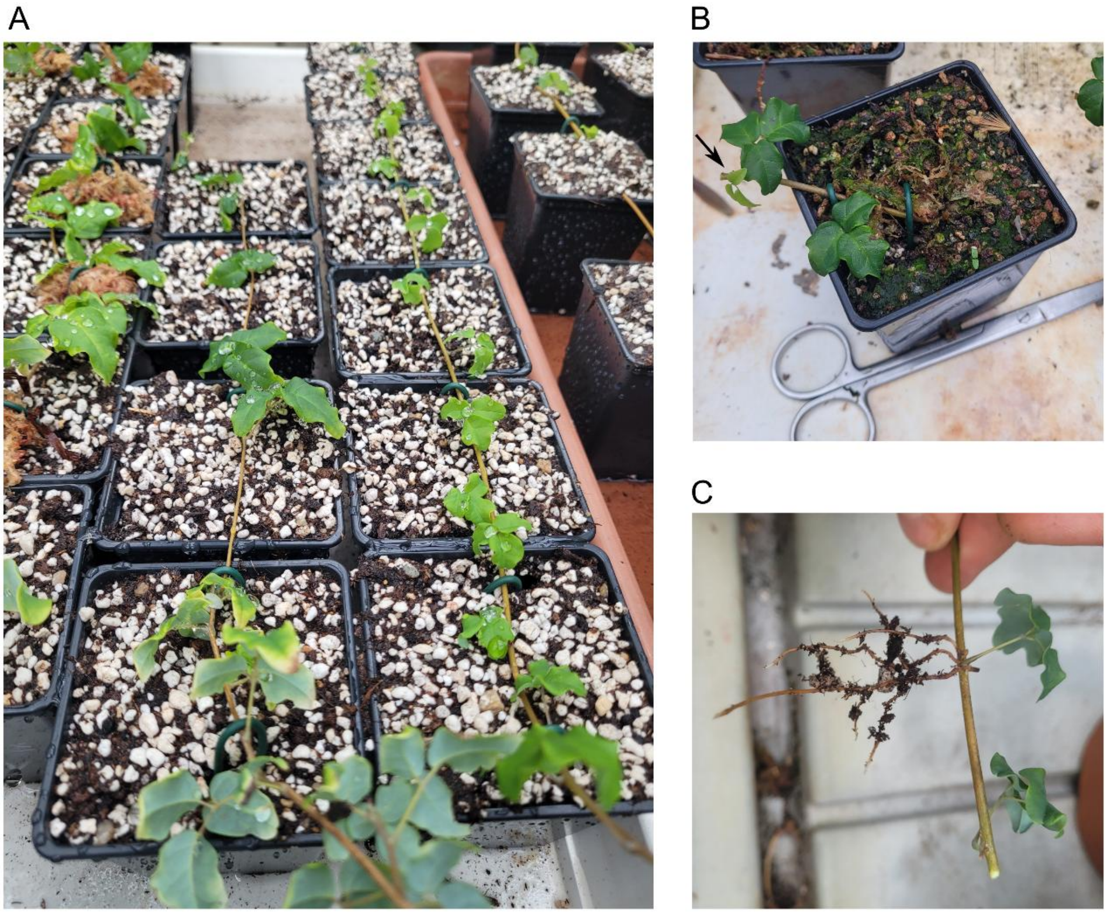


**Figure S6.** Boquila propagation by creeper shoot layering. (A) A creeper shoot attached to substrate by wire hooks. (B) A successfully rooted shoot segment after cutting of the shoot. Arrow points to a newly developing shoot. (C) Roots developed at a node of a layered segment.
